# Synchrony Genetics: Linking Ecological Mechanisms to Genetic Structure A framework for genetic inference in ecologically coupled systems

**DOI:** 10.64898/2026.04.02.716123

**Authors:** Snorre B. Hagen

## Abstract

Spatial synchrony, correlated population dynamics across space, is a defining feature of ecological dynamics, shaping outbreaks, cycles, and waves across ecosystems. Yet its genetic consequences remain poorly resolved because classical population-genetic models assume demographic independence and equilibrium conditions that synchronised populations systematically violate. Here I introduce Synchrony Genetics, a general framework that treats ecological coupling as the causal process and spatial synchrony as only one observable manifestation of that coupling. The framework links the three canonical ecological coupling mechanisms, environmental (Moran-type) coupling, dispersal-driven coupling, interaction-mediated coupling, alone or in combination, to their characteristic genetic signatures. Under this view, genetic structure is not a static property of populations or a proxy for equilibrium connectivity, but an emergent indicator of how populations are ecologically coupled across space. These expectations are synthesised in a Prediction Matrix that maps coupling mechanisms to diagnostic contrasts across widely used genetic metrics, enabling mechanism attribution from genetic data alone or in combination with demographic information. By reframing genetic patterns as evidence of coupling mechanisms rather than equilibrium processes, Synchrony Genetics provides a mechanistic foundation for interpreting genetic data in spatially coherent systems where dispersal, demographic covariance, and ecological interactions jointly shape genetic signatures. More broadly, the framework establishes a new baseline for genetic inference in systems where ecological coupling violates demographic independence, repositioning genetic structure as mechanistic evidence of how populations are linked across space.

## Introduction

Spatial synchrony, correlated (non-independent) population dynamics across space, is a pervasive feature of ecological systems. It shapes population outbreaks, cycles, and waves across ecosystems, influencing extinction risk, ecosystem stability, and the spatial footprint of ecological change (Liebhold et al. 2004). Synchronised dynamics have been documented across diverse taxa and contexts, from eruptive forest insects (Peltonen et al. 2002) and mast-seeding plants (Koenig & Knops 2013) to cyclic mammals, marine recruitment pulses, and infectious disease outbreaks (Grenfell et al. 1998). Importantly, synchrony is not restricted to outbreaking or cyclic species, but arises wherever populations are ecologically coupled. In such systems, local populations do not fluctuate independently but are linked across landscapes, often over distances far exceeding typical dispersal ranges. Here, synchrony is used in its broad ecological sense: process-level coupling that generates non-independent population trajectories across space and time. A key conceptual distinction is that ecological coupling is the causal process, whereas spatial synchrony is only one possible manifestation of that coupling.

Decades of research have established three canonical mechanisms underlying spatial synchrony (Ranta et al. 1995; Kendall et al. 2000). Shared environmental forcing synchronises populations when spatially correlated environmental variation drives parallel demographic responses (the Moran effect; Moran 1953; Lande et al. 1999). Dispersal-driven coupling arises when movement among populations transmits demographic variations across space (Bjørnstad et al. 1999; Kendall et al. 2000). Interaction-mediated synchrony emerges when shared predators, pathogens, or resources propagate variability across populations (Ims & Andreassen 2000; Post & Forchhammer 2002; Myers & Cory 2013). Although these mechanisms differ fundamentally in ecological logic, spatial scale, and causal structure, they can generate superficially similar patterns of synchronised population dynamics. As a result, demographic synchrony alone is often insufficient to identify the underlying processes coupling populations (Ranta et al. 1995).

A key reason why synchrony mechanisms are difficult to identify is that population abundance time series often collapse multiple coupling processes into similar covariance patterns (Ranta et al. 1995; Kendall et al. 2000; Bjørnstad et al. 1999; Hagen et al. 2008). Dispersal, shared environmental forcing, and trophic interactions can all generate strong synchrony, while their demographic signatures may be obscured by nonlinear responses, scale mismatches, and observational noise (Kendall et al. 2000; Liebhold et al. 2004). Consequently, the synchrony literature has repeatedly concluded that abundance data alone are often insufficient for reliable mechanism attribution (Ranta et al. 1995; Kendall et al. 2000; Liebhold et al. 2004). This limitation does not imply that synchrony mechanisms lack distinct signatures, but rather that those signatures are not always observable within a single data stream.

Genetic data provide a complementary line of evidence because they integrate realised dispersal and reproduction across generations, retaining information about coupling processes that may be invisible in demographic time series alone (Slatkin 1993; Luikart et al. 1998; Lande et al. 1999). Genetic data therefore provide leverage not by revealing synchrony itself, but by exposing the underlying coupling processes that give rise to synchronised dynamics. Despite extensive ecological understanding of synchrony, however, the genetic consequences of synchronised dynamics remain conceptually fragmented: genetic structure is commonly interpreted under equilibrium assumptions or attributed to historical contingency, rather than recognised as an emergent outcome of ongoing ecological coupling (Liebhold et al. 2004; Lande et al. 1999). This gap motivates a framework that treats genetic structure as an emergent signal of ecological coupling rather than a static population attribute, and links specific coupling mechanisms directly to their expected and diagnostic genetic signatures.

The source of this fragmentation lies in the assumption of demographic independence that underpins classical population-genetic theory. Classical models provide a clear interpretive baseline: populations are assumed to fluctuate independently, genetic drift acts locally and asynchronously, migration rates are constant through time, and allele frequencies change against a background of Hardy–Weinberg expectations (Wright 1943; Slatkin 1985, 1993; Hutchison & Templeton 1999). These assumptions are used here as an explicit baseline rather than a focus of review, because the purpose of Synchrony Genetics is to identify where and why they break down under spatially synchronised demographic coupling. Crucially, it is the ecological coupling that violates these assumptions; synchrony is merely the demographic signature of that deeper causal structure. Standard metrics such as F_ST_, isolation by distance, and clustering therefore infer connectivity under the assumption that demographic histories are uncorrelated across space. In synchronised systems, these assumptions fail: demographic fluctuations become correlated across populations, causing stochastic changes in population size and reproductive output to occur simultaneously (Lande et al. 1999; Liebhold et al. 2004). Because genetic processes are disproportionately influenced by periods of low effective population size, synchronised demographic dynamics can generate genetic signals that do not reflect local demographic histories alone (Luikart et al. 1998; Lande et al. 1999). Together, these limitations reveal the need for an approach that explicitly links ecological coupling mechanisms to their characteristic genetic consequences.

Here, I introduce Synchrony Genetics, a conceptual framework that links the ecological coupling mechanisms that generate spatial synchrony directly to their expected genetic signatures. Each synchrony mechanism imposes a distinct genetic logic, producing characteristic and diagnosable patterns in genetic data. These mechanism-specific genetic logics form the foundation of the Prediction Matrix introduced below, which links these mechanisms to diagnostic combinations of genetic differentiation, spatial structure, and clustering. Synchrony Genetics is not a population-genetic model and does not estimate demographic or dispersal parameters; instead, it provides a mechanism-aware interpretive framework for genetic patterns in ecologically coupled systems. The following sections introduce this framework in detail, outline how its predictions arise, and demonstrate its diagnostic contrasts using a minimal mechanistic simulation.

### Three ecological coupling mechanisms, three genetic logics

Spatial synchrony is not a single process but an emergent demographic outcome of distinct ecological coupling mechanisms. Each mechanism imposes characteristic constraints on movement, reproduction, and demographic coupling, producing predictable genetic signatures (Kendall et al. 2000; Liebhold et al. 2004). These contrasts are summarised schematically in Figure 1.

**Figure 1.**
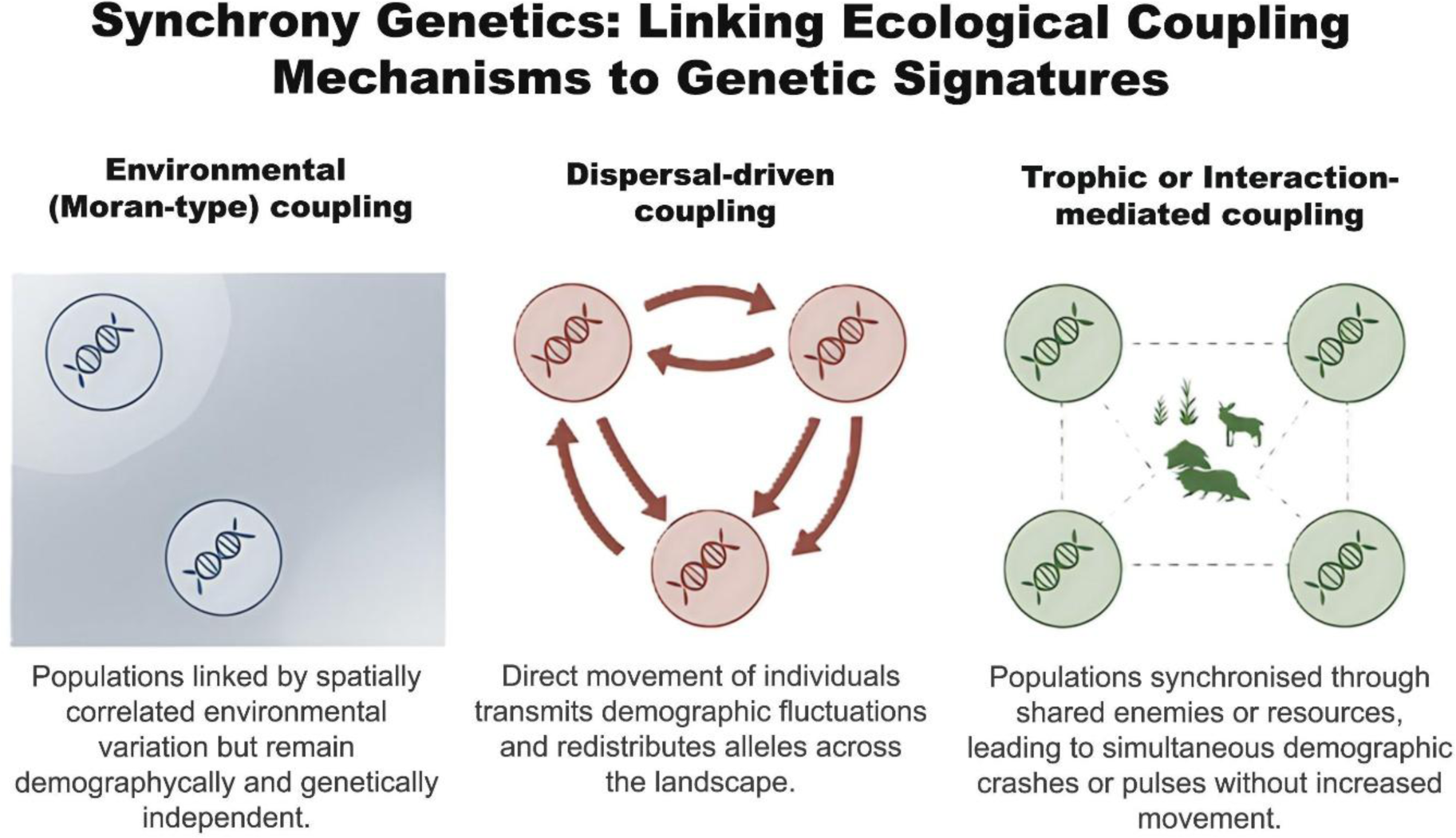
Ecological coupling mechanisms underlying spatial synchrony. Distinct ecological coupling mechanisms can generate similar spatial synchrony in population abundance, yet lead to contrasting genetic patterns. Environmental (Moran-type) coupling synchronises populations through spatially correlated environmental variation without increasing realised movement, preserving genetic independence among populations. Dispersal-driven coupling synchronises populations through direct movement of individuals, transmitting demographic fluctuations and redistributing alleles across space. Trophic or interaction-mediated coupling synchronises populations through shared enemies, pathogens, or resources, producing spatially coherent demographic impacts without increased connectivity. Natural systems may experience combinations of these mechanisms, yielding intermediate or composite genetic patterns. The figure emphasises that similar demographic synchrony can arise from distinct causal processes; interpretation therefore relies on joint genetic patterns rather than any single metric, formalised in the Prediction Matrix (Fig. 2).

### Environmental (Moran-type) coupling

Under environmental (Moran-type) coupling, populations fluctuate synchronously because they respond to spatially correlated abiotic environmental variation, such as climate anomalies or large-scale physical drivers, without requiring dispersal or biotic interactions among populations (Moran 1953; Lande et al. 1999). Populations experience similar environmental drivers, but their genetic processes remain largely independent. The genetic consequence is demographic coherence without genetic mixing. Because dispersal does not increase, allele frequencies are not homogenised across space and long-term patterns of differentiation and isolation by distance are therefore expected to persist. Parallel allele-frequency fluctuations may occur, but these reflect shared demographic forcing rather than gene flow (Liebhold et al. 2004; Hansen et al. 2020). Synchrony caused by Moran-type environmental coupling often operates at spatial scales far exceeding typical dispersal distances, explaining why populations can be tightly synchronised demographically while remaining genetically structured. This mechanism therefore provides a clear contrast to synchrony caused by dispersal-driven coupling: demography is aligned, but genes are not. In short: Moran synchrony aligns demography but not genes.

### Dispersal-driven coupling

Population synchrony caused by dispersal-driven coupling occurs when movement among populations transmits demographic fluctuations across space (Bjørnstad et al. 1999; Kendall et al. 2000). In this mechanism, synchrony requires realised movement: individuals must disperse frequently enough that local demographic variation is shared among populations. The same movements that couple population dynamics also redistribute alleles. The expected genetic signature is therefore reduced differentiation and weakened spatial structure at the scale of synchrony. When dispersal is strong enough to synchronise demography, it is typically strong enough to homogenise allele frequencies as well. Genetic clustering may be weak or absent, and isolation-by-distance relationships shallow or undetectable (Goldwyn & Hastings 2008; Abbott 2011; Anderson et al. 2018). Synchrony resulting from dispersal-driven coupling often exhibits a characteristic spatial decay: synchrony declines with distance at approximately the scale of the dispersal kernel (Ranta et al. 1995; Kendall et al. 2000; Walter et al. 2017). Here, demographic coupling and genetic homogenisation are mechanistically inseparable. Genetic structure is therefore expected to mirror this pattern, with homogenisation strongest within the synchrony domain and differentiation increasing beyond it. This mechanism therefore provides a clear contrast to Moran-type environmental coupling: demography and genes are aligned through movement. In short: dispersal-driven coupling aligns demography by mixing genes.

### Trophic or interaction-mediated coupling

Under trophic or interaction-mediated coupling, populations are synchronised through shared enemies, pathogens, or resources rather than through movement or shared abiotic forcing (Ims & Andreassen 2000; Post & Forchhammer 2002; Myers & Cory 2013). These interactions impose spatially correlated demographic change, such as predation pulses, disease outbreaks, or region-wide resource failures, without increasing realised dispersal among populations. Synchrony therefore reflects indirect ecological coupling, where populations experience similar demographic pressures at the same time despite remaining demographically and genetically isolated in space. The genetic consequences follow directly from this logic. Because dispersal does not increase, gene flow remains limited, and long-term connectivity is not strengthened. However, shared demographic stress, for example, simultaneous reductions in abundance or reproductive output, can amplify stochastic genetic change within populations concurrently across space. This can produce elevated or heterogeneous differentiation, even among populations that remain ecologically connected. Spatial structure may become patchy or weakly related to distance, because divergence reflects the timing and magnitude of shared demographic pressures rather than the geometry of dispersal. In short: trophic synchrony aligns demographic stress, not movement, and therefore can generate strong or irregular genetic structure without reducing dispersal or imposing physical barriers.

### Mixed coupling mechanisms

In natural systems, spatial synchrony rarely arises from a single ecological coupling mechanism acting in isolation. Environmental (Moran-type) coupling, dispersal-driven coupling, and trophic or interaction-mediated coupling often co-occur or operate at overlapping spatial scales, producing synchronised dynamics through multiple coupling pathways (Kendall et al. 2000; Liebhold et al. 2004; Sheppard et al. 2016). In such cases, genetic signatures are expected to reflect predictable combinations of mechanism-specific constraints rather than deviations from theory. Crucially, these hybrid genetic patterns do not represent noise, ambiguity, or failure of the framework. Instead, they are the mechanistic outcome of multiple synchronising coupling processes acting simultaneously. When coupling mechanisms operate together, each contributes its characteristic genetic influence: dispersal weakens differentiation and spatial structure, trophic interactions increase stochastic divergence through synchronised demographic impacts, and environmental forcing preserves long-term spatial patterns. Because these processes often act at different spatial scales, mixed systems can produce scale-dependent or cross-metric genetic contrasts that are diagnostic rather than problematic (Walter et al. 2017). Mixed-mechanism systems are therefore not exceptions but central cases within the framework. The Prediction Matrix accommodates these systems by identifying where intermediate, scale-dependent, or metric-discordant genetic patterns are expected. Rather than obscuring inference, mixed signatures provide additional information about the combination of ecological processes shaping synchrony, i.e. they are not noise but mechanistic information.

### From mechanism to signature: The genetic logic of Ecological Coupling and the Prediction Matrix

Ecologically coupled populations may exhibit similar synchronised abundance dynamics, yet the mechanisms that generate this synchrony impose fundamentally different constraints on movement, reproduction, and demographic correlation. As illustrated in Figure 1, these contrasts translate into distinct genetic signatures because dispersal, stochastic genetic processes, and demographic covariance interact in mechanism-specific ways (Kendall et al. 2000; Liebhold et al. 2004). Under environmental (Moran-type) coupling, synchrony arises from spatially correlated environmental variation, and genetic structure therefore reflects long-term dispersal and landscape resistance rather than demographic coherence (Moran 1953; Lande et al. 1999; Liebhold et al. 2004). Dispersal-driven coupling homogenises allele frequencies at the scale of synchrony, producing weak differentiation, shallow spatial gradients, and diffuse clustering (Bjørnstad et al. 1999; Kendall et al. 2000; Goldwyn & Hastings 2008; Abbott 2011; Anderson et al. 2018). Trophic or interaction-mediated coupling synchronises demographic impacts without increasing movement, often elevating differentiation through concurrent demographic minima and generating patchy or distance-decoupled structure (Ims & Andreassen 2000; Myers & Cory 2013; Bogdziewicz et al. 2021). Real systems frequently combine these mechanisms, producing predictable mixtures of their genetic signatures. These three genetic logics, environmental preservation, dispersal homogenisation, and interaction-driven divergence, form the conceptual backbone of the Prediction Matrix, which formalises mechanism-specific genetic contrasts into a diagnostic framework for mechanism attribution.

The Prediction Matrix synthesises these expectations using a minimal set of core population-genetic diagnostics. It summarises joint contrasts in genetic differentiation, spatial structure (isolation by distance), and clustering/admixture that distinguish environmental forcing, dispersal-driven coupling, trophic or interaction-mediated coupling, and mixed mechanisms (Figure 2). The matrix is a mechanism-centric interpretive tool: it constrains which coupling processes are compatible with an observed genetic pattern, rather than providing a unique quantitative classification or prediction. Figures 1 and 2, together with Box 1, illustrate how this diagnostic logic leads to inference by interpreting the combined genetic signal rather than any single statistic considered in isolation.

**Figure 2.**
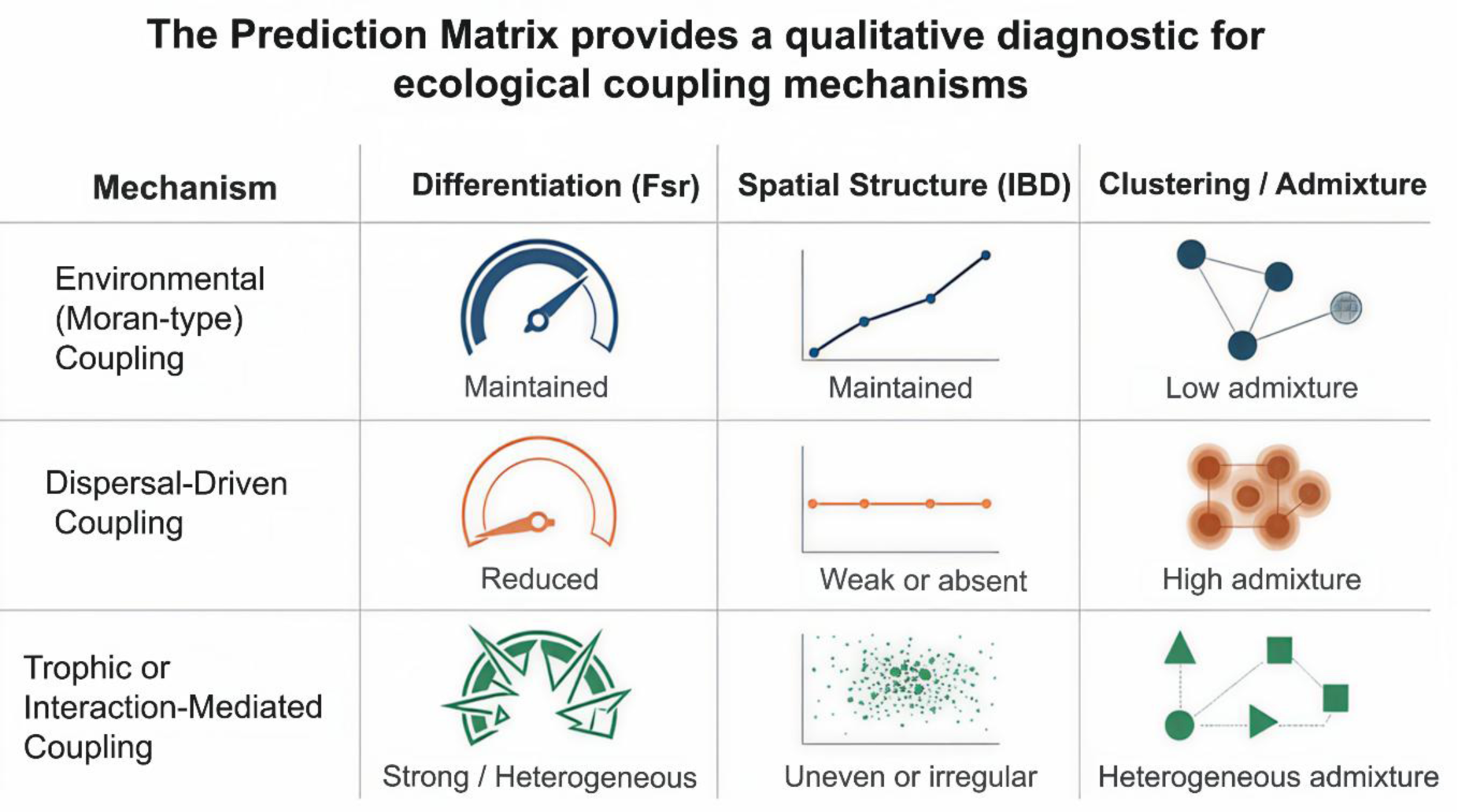
The Synchrony Genetics Prediction Matrix. Spatial synchrony can arise from distinct ecological coupling mechanisms that abundance data alone often cannot distinguish; similar demographic synchrony can therefore correspond to contrasting genetic patterns depending on the underlying coupling mechanism. The Prediction Matrix serves as a qualitative diagnostic framework that constrains which coupling processes are compatible with an observed genetic pattern, rather than providing a quantitative classifier. Inference relies on the integrated signal across metrics rather than any single statistic considered in isolation. IBD and differentiation must be interpreted relative to the spatial scale of sampling; scale-dependent patterns and mixed mechanisms are expected and are summarised in Supplementary Table S1, which provides a complete tabular version and operational guidance for interpretation.

#### Box 1. Diagnosing ecological coupling mechanisms from genetic patterns

**Purpose.** The Prediction Matrix provides a diagnostic framework for interpreting genetic structure in ecologically coupled systems, with spatial synchrony as one prominent example.

**Scenario.** A species exhibits spatially coherent demographic fluctuations across a region. Genetic data are available from multiple populations within the domain of ecological coupling (the spatial extent over which abundance dynamics are strongly correlated).

**Pattern A —** Structure maintained under ecological coupling**. Genetic pattern:** Persistent spatial structure; isolation-by-distance present or unchanged; little or no admixture. **Interpretation:** Environmental (Moran-type) coupling. Spatially correlated environmental variation synchronises demography without increasing realised connectivity. Genetic structure therefore reflects long-term dispersal processes rather than the demographic coherence itself.

**Pattern B —** Structure eroded at the coupling scale**. Genetic pattern:** Weak or absent differentiation; indistinct clustering and/or widespread admixture; shallow or absent IBD. **Interpretation:** Dispersal-driven coupling. Movement transmits demographic variation across space and simultaneously homogenises allele frequencies. Demographic coherence and genetic mixing are mechanistically linked.

**Pattern C —** Strong or heterogeneous structure without admixture. **Genetic pattern:** Strong or patchy differentiation; limited admixture; spatial structure weakly related to geographic distance**. Interpretation:** Trophic or interaction-mediated coupling. Shared predators, pathogens, or resources impose spatially coherent demographic impacts without increased dispersal. Genetic structure reflects the timing and magnitude of shared demographic pressures rather than dispersal-driven mixing.

**Key point.** Similar demographic coherence can arise from distinct ecological mechanisms. The joint genetic pattern therefore acts as a mechanism filter, constraining which coupling processes are compatible with the observed data and reframing genetic structure as evidence of ecological coupling rather than a static population attribute. More broadly, the diagnostic logic illustrated here shows how Synchrony Genetics can be generalised toward a broader Coupling Genetics framework that interprets genetic structure through the lens of ecological coupling rather than equilibrium assumptions.

### Scale-Dependence of Genetic Signatures

The genetic expectations in the Prediction Matrix are inherently scale-dependent. Isolation-by-distance (IBD), differentiation, and clustering all emerge from the interaction between dispersal distance and the spatial extent over which populations are sampled (Wright 1943; Slatkin 1993; Hutchison & Templeton 1999). Because all biological populations experience some degree of movement, spatial genetic patterns always reflect the interplay between mechanism-specific processes and the broader dispersal landscape rather than any single process acting in isolation (Slatkin 1993; Broquet & Petit 2009).

Under Moran-type environmental coupling, synchrony does not increase movement, and genetic patterns therefore reflect the baseline spatial structure, which may be weak or strong depending on the spatial scale of sampling relative to background dispersal. Moran forcing preserves the underlying genetic landscape rather than generating new patterns, consistent with theory predicting demographic synchrony without increased gene flow under spatially correlated environmental variation (Moran 1953; Lande et al. 1999; Liebhold et al. 2004).

In contrast, dispersal-driven coupling produces synchrony and homogenises allele frequencies within the synchrony domain (the spatial extent over which abundance dynamics are strongly correlated). When the effective scale of movement exceeds the sampling extent, differentiation and IBD collapse; when sampling extends beyond this domain, IBD re-emerges. Movement over short distances can still produce gradual spatial changes in allele frequencies across large regions, whereas movement over longer distances mixes populations much more rapidly, yielding scale-dependent transitions in genetic structure (Slatkin 1985, 1993; Hutchison & Templeton 1999). Thus, the prediction of “weak or absent IBD” applies within, but not necessarily beyond, the spatial scale at which dispersal couples demographic dynamics, consistent with synchrony theory linking dispersal kernels to spatial decay of coherence (Ranta et al. 1995; Kendall et al. 2000; Walter et al. 2017).

Trophic or interaction-mediated coupling can generate spatially coherent demographic erosion without increasing movement, producing patchy or irregular structure whose spatial expression depends on the scale and synchrony of demographic minima rather than the geometry of dispersal. Such patterns reflect synchronised demographic impacts that restructure genetic variation independently of dispersal geometry (Lande et al. 1999; Luikart & Cornuet 1998; Ims & Andreassen 2000).

Mixed coupling mechanisms illustrate these principles clearly: dispersal may weaken differentiation at short distances while environmental forcing or trophic interactions shape divergence at broader spatial scales. Such systems naturally produce composite, scale-specific genetic signatures, reflecting the superposition of multiple coupling processes operating at different spatial extents rather than ambiguity or failure of inference (Kendall et al. 2000; Liebhold et al. 2004; Walter et al. 2017).

Recognising these scale dependencies is essential for correct application of Synchrony Genetics, and, more generally, for interpreting genetic structure under ecological coupling: the diagnostic logic of the framework rests not on absolute values of F_ST_ or IBD, but on how their spatial patterns, and their scale transitions, align with the expected mechanism-specific combinations (Wright 1943; Slatkin 1993; Hutchison & Templeton 1999). In practice, Synchrony Genetics should be applied by (i) matching the spatial scale of genetic sampling to the observed synchrony scale, and (ii) explicitly checking whether diagnostic patterns change when the spatial window is expanded or restricted. Thus, the Prediction Matrix should be interpreted at the spatial scale where synchrony is expressed, not necessarily across the entire sampling extent. Scale dependence does not weaken the Prediction Matrix; it is built into its logic.

### Simulation of Mechanism-Specific Genetic Signatures

To illustrate the diagnostic patterns summarised in the Prediction Matrix, we conducted a minimal forward-time simulation (Figure 3). Details of the simulation model and all parameter settings are provided in Supplementary Note S1. The results show that the genetic signatures predicted by Synchrony Genetics arise directly from the ecological coupling mechanisms that generate synchronised dynamics, rather than from any assumed synchrony or imposed genetic effects, consistent with theory linking dispersal, environmental forcing, and demographic correlation to synchrony (Kendall et al. 2000; Liebhold et al. 2004). The simulation therefore serves as a mechanistic sanity check: the diagnostic contrasts in the Prediction Matrix emerge directly from ecological coupling rules, without tuning genetic parameters to force the outcome.

**Figure 3.**
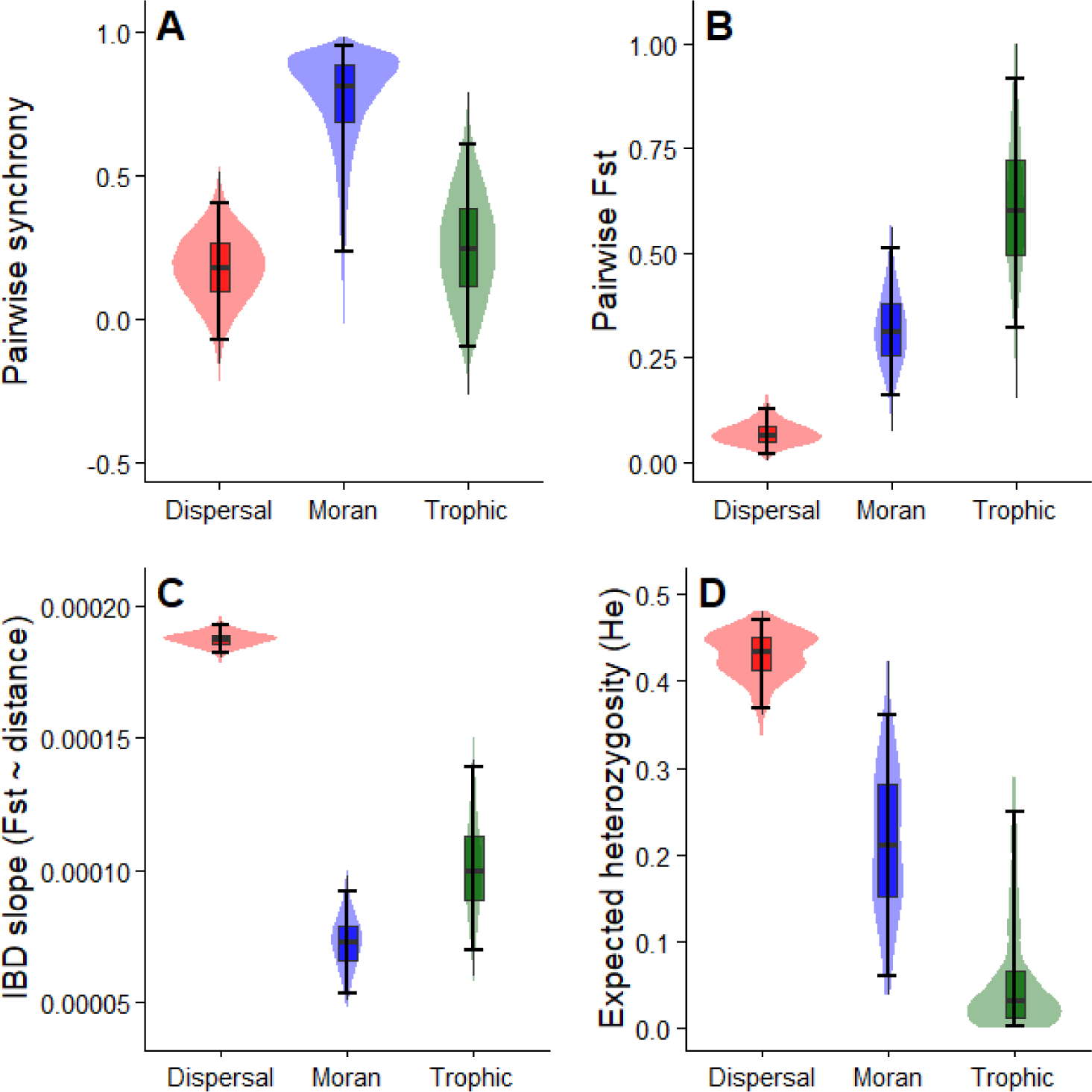
Mechanistic simulation of spatial genetic signatures under alternative synchrony mechanisms. A minimal forward-time simulation (300 demes) illustrates how distinct ecological coupling processes generate characteristic spatial genetic patterns. A. Pairwise synchrony: Environmental (Moran-type) coupling produces moderate synchrony without gene flow; dispersal-driven coupling yields strong demographic coherence; trophic or interaction-mediated coupling generate highly variable synchrony. B. Pairwise F_ST_: Differentiation remains low under Moran-type environmental coupling, is reduced within the synchrony domain under dispersal-driven coupling, and is elevated and heterogeneous under trophic coupling due to synchronised demographic erosion. C. Isolation-by-distance (IBD) slopes: Moran-type environmental coupling maintains a clear positive IBD signal. Dispersal-driven coupling reduces differentiation locally, but short-range movement can generate gradual allele-frequency changes across the full landscape before global homogenisation is complete. Trophic coupling produces irregular or intermediate IBD patterns consistent with synchronised demographic impacts on genetic structure. D. Expected heterozygosity (H_e_): Heterozygosity is highest under dispersal-driven coupling and lowest under trophic coupling. Together, these patterns provide a mechanistic demonstration of the diagnostic contrasts summarised in the Prediction Matrix. A high-resolution version of this figure is provided as Supplementary Figure S1.

Across a landscape of 300 local populations, Moran environmental forcing produced low but spatially structured genetic differentiation with a clear positive isolation-by-distance (IBD) signal. This pattern reflects the preservation of the baseline spatial structure generated by limited background dispersal. Because Moran-type environmental coupling does not increase movement, it maintains rather than alters this underlying genetic landscape, as expected under synchrony driven by spatially correlated environmental variation rather than gene flow (Moran 1953; Lande et al. 1999; Liebhold et al. 2004).

Dispersal-driven coupling generated high demographic coherence together with markedly reduced differentiation within the synchrony domain. Across the full landscape, however, differentiation declined more slowly than under Moran-type environmental coupling. This reflects the short-range movement used in the simulation: frequent exchanges among neighbouring populations rapidly homogenise allele frequencies locally, but it takes longer for these changes to spread across the entire region. As a result, smooth spatial gradients can appear temporarily even when dispersal is the mechanism generating synchrony, a behaviour consistent with classic expectations for spatial genetic change under limited dispersal (Slatkin 1993; Hutchison & Templeton 1999). This is a consequence of dispersal scale relative to landscape size, not a contradiction of dispersal-driven coupling. The key point is that dispersal-driven synchrony refers to the ecological mechanism rather than any specific movement pattern, consistent with theory showing that synchrony depends on realised dispersal rather than the specific pattern of movement (Bjørnstad et al. 1999; Kendall et al. 2000; Goldwyn & Hastings 2008). If individuals occasionally moved longer distances, homogenisation would occur rapidly across the whole landscape and spatial structure would collapse everywhere, producing the canonical genetic signature of dispersal-driven coupling (Slatkin 1985; Broquet & Petit 2009). The contrast between short- and long-distance movement therefore reflects differences in dispersal scale, not differences in ecological mechanism.

Trophic or interaction-mediated coupling produced the strongest and most heterogeneous divergence and substantially reduced heterozygosity, reflecting spatially coherent demographic erosion rather than increased movement. Because synchronised demographic crashes impose strong, spatially aligned demographic constraints, the resulting genetic structure reflects the scale and synchrony of demographic impacts rather than dispersal geometry, consistent with theory linking demographic bottlenecks to elevated differentiation (Lande et al. 1999; Luikart & Cornuet 1998). This yields patchy or irregular spatial patterns, in line with expectations for interaction-mediated synchrony driven by shared enemies or resources rather than dispersal (Ims & Andreassen 2000; Myers & Cory 2013).

Recognising these scale dependencies clarifies why the relative ordering of mechanisms in the simulation matches the Prediction Matrix, even though the absolute magnitude of IBD differs across kernel shapes and spatial scales. Synchrony therefore arises mechanistically from each coupling process in every scenario; it is never imposed. Genetic patterns emerge solely from how each coupling mechanism structures movement, reproduction, and demographic variation, reinforcing the distinction between ecological coupling as a causal process and synchrony as one statistical manifestation of that coupling (Ranta et al. 1995; Kendall et al. 2000; Liebhold et al. 2004). This distinction is central to generalising Synchrony Genetics into a broader Coupling Genetics perspective.

### Temporal considerations and scope of the framework

Genetic data integrate demographic and dispersal processes across generations, and genetic structure therefore reflects not only how populations are ecologically coupled, but also when, for how long, and with what temporal pattern that coupling operates (Slatkin 1993; Lande et al. 1999). In synchronised systems, ecological coupling may be persistent, episodic, or pulsed, and genetic signatures may track coupling closely, lag behind it, or persist long after coupling has weakened or disappeared, reflecting non-equilibrium dynamics and genetic memory (Slatkin 1993; Luikart & Cornuet 1998).

The framework developed here is intentionally static. Its purpose is to isolate mechanism-specific genetic contrasts under spatial synchrony, rather than to describe the temporal evolution of genetic patterns explicitly. Establishing these contrasts is a necessary first step, because temporal genetic dynamics depend on the underlying synchrony mechanism: dispersal-mediated coupling, environmental forcing, and interaction-mediated coupling impose fundamentally different constraints on the timing and strength of genetic changes (Kendall et al. 2000; Liebhold et al. 2004).

Explicit treatment of temporal dynamics, including genetic response times, lag effects, and genetic memory under changing coupling regimes, represents a substantial extension of the framework and lies beyond the scope of the present manuscript. Importantly, incorporating temporal dynamics does not weaken the static diagnostics developed here; rather, it builds directly upon them. Mechanism-specific contrasts provide the essential reference against which temporal change can be interpreted, ensuring that transient or legacy genetic signals are not misattributed to ongoing ecological processes (Slatkin 1993; Luikart & Cornuet 1998). Temporal dynamics modify the expression of these patterns but not the underlying mechanism-specific contrasts.

Modern genomic datasets fit naturally within this framework because Synchrony Genetics is mechanism-centric rather than marker-centric. High-resolution genomic data do not change the underlying logic of the Prediction Matrix; instead, they increase power to detect the joint patterns of differentiation, spatial structure, and admixture that diagnose coupling mechanisms. Dense SNP datasets improve estimation of F_ST_ and isolation-by-distance slopes, while whole-genome data enhance inference of drift intensity, effective population size, and temporal allele-frequency covariance (Hutchison & Templeton 1999; Broquet & Petit 2009). Importantly, genomic resolution does not erase mechanism-specific contrasts: dispersal-driven coupling still produces homogenisation, Moran-type coupling still preserves long-term structure, and trophic or interaction-mediated coupling still generates strong or heterogeneous differentiation. Genomic data therefore strengthen, rather than alter, the diagnostic capacity of the framework.

I refer to this framework as Synchrony Genetics because spatial synchrony is the most visible and widely studied expression of ecological coupling (Liebhold et al. 2004; Walter et al. 2017). However, the underlying logic is more general: genetic data can diagnose the ecological coupling processes that generate non-independent demographic trajectories, whether or not synchrony is the dominant manifestation. In this broader sense, the framework can be viewed as a form of Coupling Genetics, with synchrony representing one prominent case within a wider class of coupling architectures.

Ecological coupling is broader than spatial synchrony, and many coupling processes link populations without producing correlated abundance dynamics. Such processes can also restructure genetic landscapes, but they do so through pathways that do not manifest as synchrony and therefore fall outside the scope of the present framework. Synchrony is used here as a tractable and well-characterised case because ecological coupling is directly observable and because synchrony generates spatial autocorrelation in demographic rates, pushing genetic interpretation into territory where independent population assumptions break down (Ranta et al. 1995; Liebhold et al. 2004). The coupling-centric logic developed here is general, however, and provides a foundation for future extensions to non-synchronising forms of ecological coupling (Box 2).

#### Box 2. Ecological coupling extends beyond synchrony

**Purpose.** Ecological coupling occurs when populations share ecological processes that make their demographic histories partially non-independent. Spatial synchrony is only one expression of such coupling. Many coupling processes link populations without producing correlated abundance dynamics, yet still reshape genetic structure. This box highlights how the coupling-centric logic of Synchrony Genetics generalises beyond synchronising systems.

**Examples of ecological coupling.** Populations may be coupled through asymmetric source–sink dynamics, density-dependent dispersal, shared predators or mutualists, behavioural interactions such as conspecific attraction, reproductive interference, or landscape-mediated demographic dependence. These processes alter movement, reproduction, or demographic sensitivity across space, creating non-independent population trajectories even when abundance time series remain uncorrelated.

**Genetic implications.** Non-synchronising coupling can generate directional gene flow, asymmetric differentiation, spatially structured admixture, or context-dependent clustering. These patterns arise from the mechanisms of coupling rather than from synchrony itself. Genetic structure integrates these shared demographic histories over generations, reflecting how populations are ecologically linked rather than how they fluctuate.

**Key point.** Synchrony provides a tractable case where ecological coupling is directly observable, allowing mechanism-specific genetic contrasts to be established clearly. The coupling-centric logic is broader, however, and forms a foundation for future extensions to non-synchronising forms of ecological coupling.

### What synchrony genetics enables

Synchrony Genetics shifts genetic interpretation from description to mechanism in ecologically coupled systems. Rather than asking whether populations are genetically structured, the framework asks why particular patterns arise, given how populations are ecologically coupled (Liebhold et al. 2004). Genetic structure is thus treated not as a static population attribute, but as an emergent ecological signal shaped by the mechanisms that synchronise population dynamics.

The ecological mechanisms that couple populations also leave signatures in demographic time series. However, as discussed above, dispersal, shared environmental forcing, and trophic interactions frequently collapse into similar covariance patterns in abundance data, limiting their power for reliable mechanism attribution (Ranta et al. 1995; Kendall et al. 2000; Bjørnstad et al. 1999; Liebhold et al. 2004). Genetic data provide a complementary line of evidence because they integrate realised dispersal, reproduction, and demographic history across generations, thereby retaining mechanistic information that may be obscured by nonlinear responses, scale mismatches, and observational noise in demographic dynamics (Luikart et al. 1998; Lande et al. 1999; Slatkin 1993).

A central capability of the framework is mechanism attribution. In many systems, demographic time series alone cannot distinguish whether coupling arises from shared environmental forcing, dispersal, trophic interactions, or combinations thereof (Kendall et al. 2000). Because genetic data integrate realised dispersal and reproduction across generations, they retain information about coupling processes that may be invisible in abundance data alone. When interpreted through Synchrony Genetics, joint patterns of genetic differentiation, isolation by distance, and clustering can therefore be used to distinguish movement-driven coupling from environmentally forced or interaction-mediated coupling.

By focusing on mechanisms rather than taxa or life histories, Synchrony Genetics facilitates synthesis across systems. The framework applies equally to insects, plants, vertebrates, marine organisms, and pathogens (Peltonen et al. 2002; Gouveia et al. 2016), providing a common interpretive language for systems that differ widely in ecology, scale, and life history. This generality becomes increasingly valuable as spatial synchrony intensifies across ecosystems under global change (Walter et al. 2017; Hansen et al. 2020).

Finally, Synchrony Genetics provides a conceptual bridge between ecological and evolutionary processes by shifting genetic interpretation from pattern description to process attribution. By explicitly linking ecological coupling mechanisms to their characteristic genetic signatures, the framework clarifies when genetic structure reflects ongoing movement and when it reflects longer-term landscape constraints imposed by ecological coupling. This integration enables comparative analyses across systems, supports synthesis across spatial and temporal scales, and establishes a foundation for predictive approaches that jointly consider ecological coupling and evolutionary consequences.

### Outlook

The framework developed here has immediate consequences for how genetic data are interpreted in ecologically coupled systems, of which spatial synchrony represents a prominent and well-studied case (Liebhold et al. 2004). By explicitly linking ecological coupling mechanisms to genetic expectations, Synchrony Genetics replaces descriptive pattern matching with mechanistic inference under nonequilibrium ecological conditions. Spatial synchrony is treated here as a canonical and empirically tractable manifestation of ecological coupling, allowing general principles linking ecological coupling to genetic structure to be developed within a well-established theoretical context. Once ecological coupling is recognised as a primary determinant of genetic structure, considerations of history, time, and mechanism follow directly from the logic of Synchrony Genetics. Because coupling mechanisms generate non-independent demographic and genetic histories across populations (Lande et al. 1999; Liebhold et al. 2004), equilibrium interpretations become increasingly fragile as coupling strengthens.

Ecological coupling therefore forces explicit consideration of time. Genetic patterns integrate demographic and dispersal processes across generations, and their interpretation depends on when, for how long, and at what spatial scale populations are coupled. Under persistent coupling, genetic structure may track coupling mechanisms closely; under episodic or transient coupling, genetic signatures may lag behind, persist as historical imprints, or mislead if interpreted without mechanistic context (Lande et al. 1999). These temporal consequences arise inherently from ecological coupling and do not require additional assumptions beyond those established here. Distinguishing coupling from synchrony also clarifies that the same logic applies in time: temporal genetic patterns reflect the timing, duration, and magnitude of ecological coupling, not temporal synchrony per se.

Finally, synchrony genetics carries clear implications for inference and monitoring. Because ecological coupling produces correlated demographic and genetic processes across space, reliance on single metrics or equilibrium expectations becomes increasingly unreliable (Kendall et al. 2000; Liebhold et al. 2004). Interpreting genetic data in ecologically coupled systems therefore requires integrative, mechanism-aware diagnostics that consider multiple genetic summaries jointly. This need becomes especially acute as environmental forcing, connectivity, and trophic interactions increasingly align population dynamics under global change (Hansen et al. 2020).

Together, these considerations position Synchrony Genetics not only as a diagnostic framework, but as a foundation for theory, empirical synthesis, and applied monitoring. By explicitly linking ecological coupling to evolutionary consequences, the framework opens a path toward a more dynamic and predictive understanding of how populations are connected across space and time. Key next steps include extending the framework to explicit temporal dynamics, integrating it with landscape-genomic models, and applying it to long-term monitoring programmes where ecological coupling is intensifying under global change.

### Conclusion

Ecological coupling fundamentally alters what genetic structure means. In spatially synchronised systems, one prominent manifestation of ecological coupling, populations are not independent demographic units, and genetic patterns cannot be interpreted as equilibrium connectivity by default (Lande et al. 1999; Liebhold et al. 2004). Synchrony Genetics reframes population genetics as mechanistic inference: rather than asking whether populations are genetically structured, it asks why structure takes the form it does, given how populations are ecologically coupled across landscapes.

In ecologically coupled systems, unexplained or discordant genetic patterns should not be dismissed as noise or equilibrium artefacts, but recognised as informative outcomes of correlated demography and coupling (Liebhold et al. 2004; Kendall et al. 2000). Genetic structure thus functions as an ecological signal, shaped jointly by dispersal and correlated demographic processes, rather than as a static population trait retrofitted to equilibrium narratives.

By explicitly linking the canonical ecological coupling mechanisms, Moran-type environmental coupling, dispersal-driven coupling, and trophic or interaction-mediated coupling, to diagnostic contrasts across widely used genetic metrics, the framework converts common genetic summaries into evidence about ecological mechanisms. The Prediction Matrix formalises this logic and clarifies why similar demographic coherence can correspond to opposing genetic outcomes (Kendall et al. 2000; Liebhold et al. 2004).

Synchrony Genetics therefore shifts the role of genetic data from descriptive patterns to mechanistic evidence of how populations are connected across space. In ecologically coupled systems, interpreting genetic structure without reference to coupling mechanisms is no longer a conservative default. The framework establishes a new baseline for interpreting genetic data in ecologically coupled systems, replacing equilibrium connectivity as the default lens for interpreting genetic patterns.

## Acknowledgements

I thank my colleagues at NIBIO, Hans Geir Eiken and Paul Eric Aspholm, for reading the manuscript and providing constructive feedback and discussions that helped refine the ideas presented here and improve clarity and presentation.

## Author contributions

S.B.H. conceived the study, developed the conceptual framework, and wrote the manuscript.

## Data availability

This study did not generate or analyse empirical data. All simulation output is reproducible from the code provided in the Supplementary Information.

## Code availability

All simulation code used in this study is provided in full in Supplementary Note S2 and can be run directly to reproduce all results. The simulations were conducted using R version 4.3.1 (R Core Team 2023).

## SUPPLEMENTARY INFORMATION

### Supplementary Note S1 — Simulation Methods

#### S1.1 Purpose

This simulation illustrates how distinct ecological coupling mechanisms, Moran environmental forcing, dispersal-driven synchrony, and trophic (interaction-mediated) synchrony, generate different spatial genetic signatures. The goal is not to model any specific species, but to demonstrate mechanistic links between coupling processes and genetic outcomes under minimal assumptions. Synchrony is never imposed; it emerges from ecological processes. Genetic patterns arise from the demographic trajectories generated by each mechanism.

#### S1.2 Model overview

We implemented a forward-time individual-based simulation across:

- **300 demes** arranged in a 1-D ring
- **100 individuals per deme**
- **20 biallelic loci**
- **300 generations**, last 100 used for diagnostics
- **Neutral genetic processes:** mutation, reproduction, and limited dispersal

A common simulation structure was used for all mechanisms; only the ecological coupling rules differ.

### S1.3 Synchrony mechanisms

#### Environmental (Moran-type) environmental coupling

Spatially correlated environmental noise generates demographic synchrony without increasing dispersal or altering long-term connectivity.

#### Dispersal-driven coupling

High dispersal (m = 0.90) transmits local demographic variation across demes and homogenises allele frequencies.

#### Trophic or interaction-mediated coupling

Periodic synchronous demographic crashes (every 5 generations, 15% survival) impose strong, spatially aligned demographic constraints across demes, creating patchy or strong genetic divergence.

### S1.4 Parameters

| Parameter | Value | Notes |
| --- | --- | --- |
| Demes | 300 | 1D ring |
| Individuals per deme | 100 | Constant N |
| Loci | 20 | Neutral biallelic |
| Generations | 300 | 200 burn-in + 100 analysis |
| Mutation rate | $1 \times 10^{-3}$ | Symmetric |
| Dispersal (Moran, Trophic) | 0.01 | Low |
| Dispersal (Dispersal-driven) | 0.90 | High |
| Crash interval | 5 | Trophic only |
| Crash survival | 0.15 | Trophic only |
| Environmental noise | scenario-specific | Moran vs others |

### S1.5 Diagnostics computed

- Pairwise synchrony: correlation of standardised abundance series
- Pairwise: F_ST_ standard H_t_ - H_s_ estimator
- Expected heterozygosity (H_e_): mean per-deme gene diversity
- Isolation-by-distance (IBD) slope: linear regression of F_ST_ ring distance; bootstrapped CIs

### S1.6 Summary of results

**Table S1.** Mean values and 95% confidence intervals for all four genetic diagnostics across the three ecological coupling mechanisms generating synchrony (300 demes). Values are derived from the forward-time simulation described in Supplementary Note S1 and calculated from the full pairwise matrices (synchrony, FST, IBD, He) and bootstrap distributions (IBD slopes).

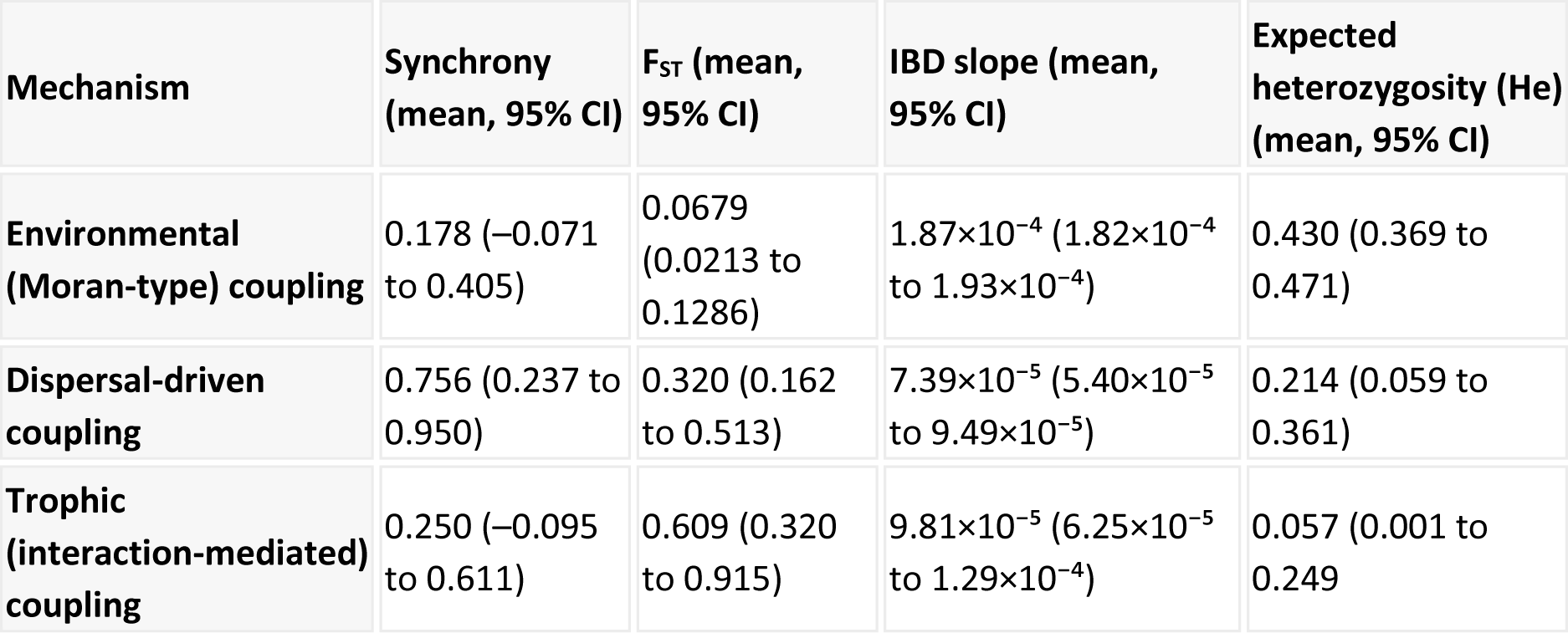

These align with the diagnostic patterns in the Prediction Matrix:

- Environmental (Moran-type) coupling → differentiation maintained, IBD strong
- Dispersal-driven coupling → homogenisation, IBD weak
- Trophic or interaction-mediated coupling → strong/heterogeneous divergence, low He

### S1.7 Supplementary Figure S1

**Figure S1.**
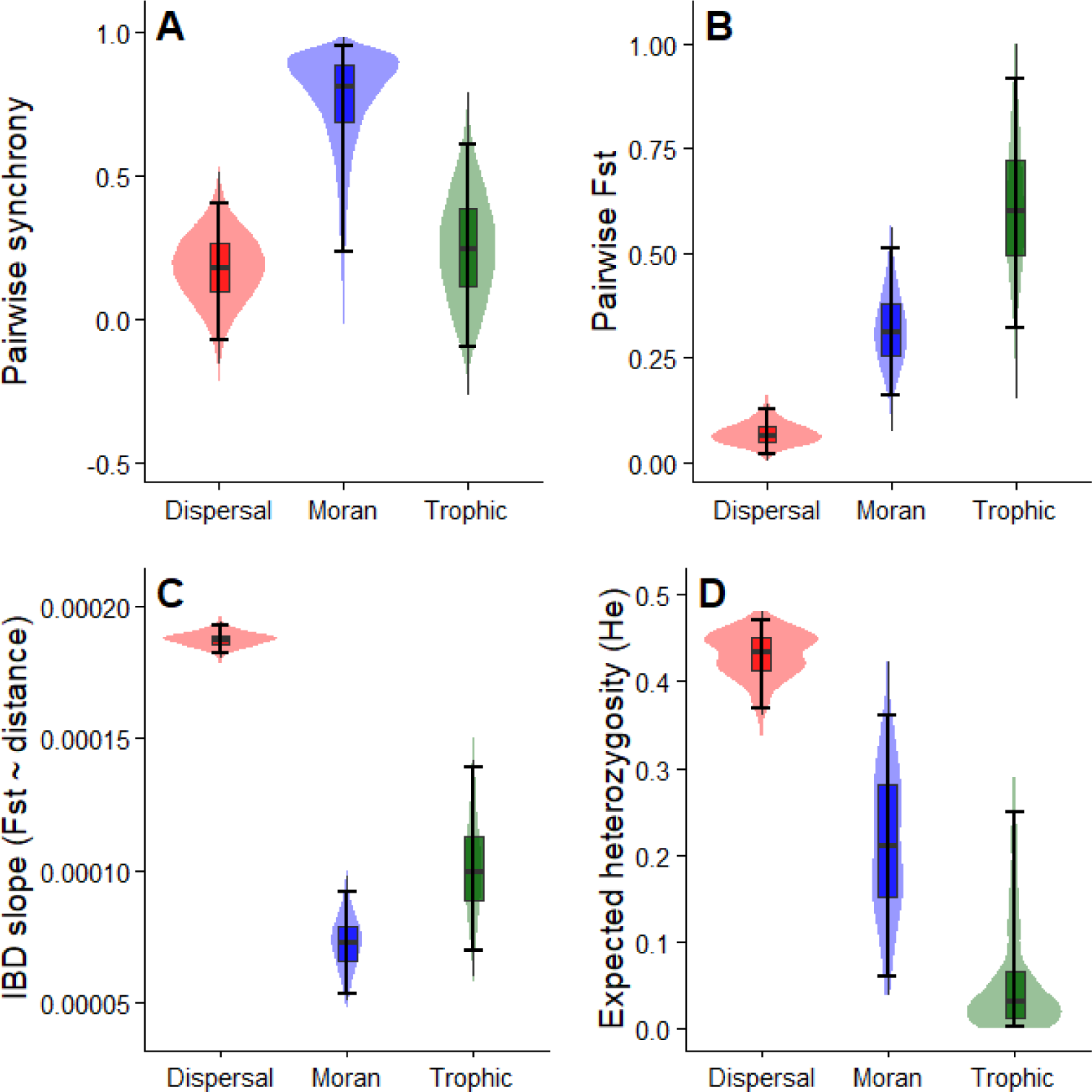
Full-resolution simulation of mechanism-specific spatial genetic signatures. High-resolution version of Figure 2. Violin distributions, medians, and 95% confidence intervals are shown for all genetic metrics across 300 demes. A. Pairwise synchrony: Moran forcing generates moderate synchrony driven by shared environmental noise; dispersal-driven synchrony yields the highest coherence across demes; trophic synchrony exhibits wide variation due to intermittent, synchronised demographic crashes. B. Pairwise F_ST_: Moran forcing maintains low but structured differentiation; dispersal-driven synchrony homogenises allele frequencies; trophic synchrony produces both the strongest and most heterogeneous F_ST_ values (ranging up to ∼0.9). C. IBD slopes: Moran forcing retains a stable positive IBD pattern, dispersal-driven synchrony collapses IBD (approaching zero), and trophic coupling yields intermediate slopes reflecting patchy structure generated by synchronised demographic impacts. D. Expected heterozygosity (H_e_): H_e_ is highest under dispersal-driven synchrony, intermediate under Moran forcing, and lowest under trophic coupling due to spatially coherent demographic erosion. Together, these diagnostics provide a mechanistic demonstration that the ecological processes producing synchrony also generate distinct and predictable genetic signatures, consistent with the Prediction Matrix.

### S1.8. Supplementary Table S1

**Table S1.** Prediction Matrix linking ecological coupling mechanisms to diagnostic genetic patterns. This table provides the full tabular version of the Prediction Matrix summarised graphically in Fig. 2. It lists the expected joint patterns of genetic differentiation, spatial structure (isolation by distance), and clustering/admixture under alternative ecological coupling mechanisms that can generate spatial synchrony, including environmental (Moran-type) coupling, dispersal-driven coupling, trophic or interaction-mediated coupling, and mixed coupling mechanisms. The matrix is intended as a qualitative diagnostic reference for mechanism attribution. Inference should be based on the joint pattern across metrics rather than any single statistic considered in isolation. Scale-dependent effects and mixed coupling regimes are expected and are explicitly accommodated.

| Mechanism | Genetic differentiation | Spatial structure / isolation by distance | Clustering / admixture | Diagnostic cue |
| --- | --- | --- | --- | --- |
| **Moran forcing (shared environment)** | Differentiation maintained | Baseline spatial structure and IBD maintained | Distinct clusters; low admixture | Synchrony without mixing; structure reflects long-term connectivity |
| **Dispersal-driven synchrony** | Differentiation reduced within the synchrony domain | Spatial structure and IBD weak or absent within the synchrony domain | Diffuse clustering; high admixture | Synchrony with gene flow; homogenisation at synchrony scale |
| **Trophic or interaction-mediated synchrony** | Strong or heterogeneous differentiation | Uneven or irregular spatial structure | Distinct clusters; inconsistent admixture patterns | Synchronous demographic impacts generate pronounced or heterogeneous structure |
| **Mixed mechanisms** | Intermediate or context-dependent differentiation | Scale-dependent spatial structure | Metrics may diverge | Combined effects of multiple mechanisms |

**How to use this table.** Interpretation should be based on the joint pattern across differentiation, IBD, and clustering, rather than any single metric. The appearance of these patterns can vary with spatial scale, so the contrasts in the table refer to the scale at which synchrony is generated relative to sampling extent and background dispersal; further details on scale.dependence are provided in the following section. Apparent mismatches among metrics may indicate mixed mechanisms or sampling limitations.

### Supplementary Note S2 — Complete R Script for Reproducibility

The script below can be copied and run directly in R without modification. It reproduces the entire simulation workflow used in this study, including: (1) generating demographic time series under the three ecological coupling mechanisms underlying spatial synchrony; (2) simulating genotypes across 300 demes through 300 generations; (3) computing all diagnostic metrics (pairwise synchrony, pairwise F_ST_, IBD slopes, H_e_); and (4) producing all summary estimates, confidence intervals, and the mechanistic simulation figure shown in the main text and Supplementary Figure S1. This script was developed and executed using R version 4.3.1 (R Core Team 2023).

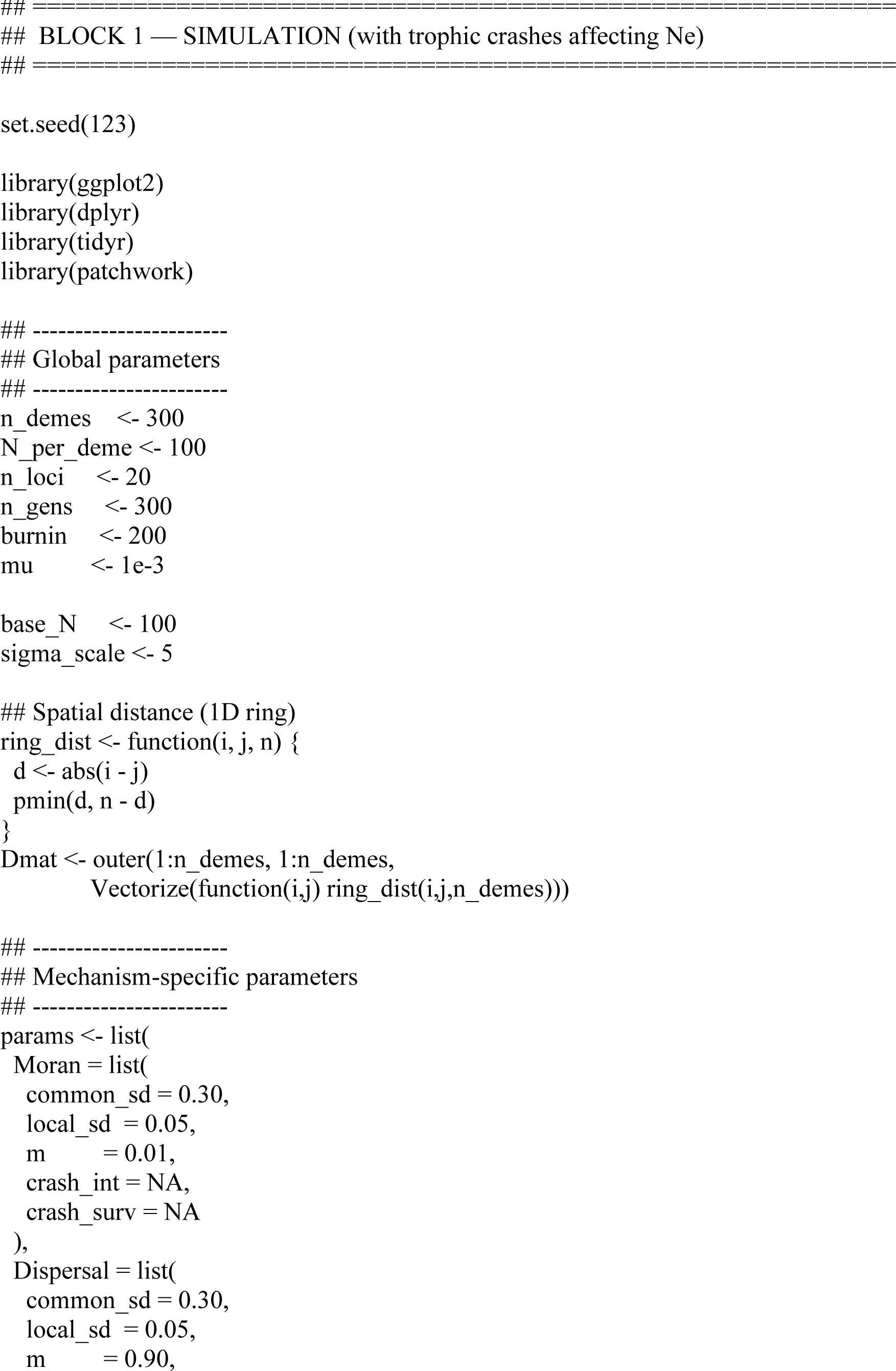
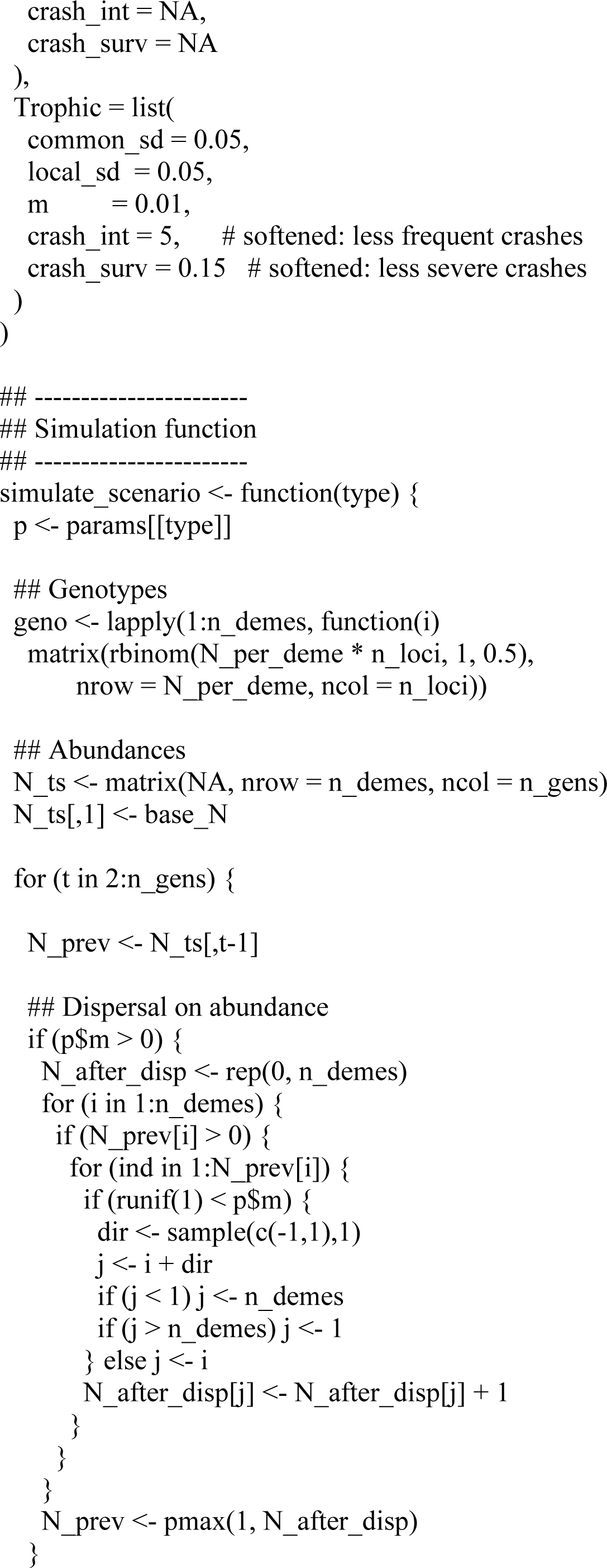
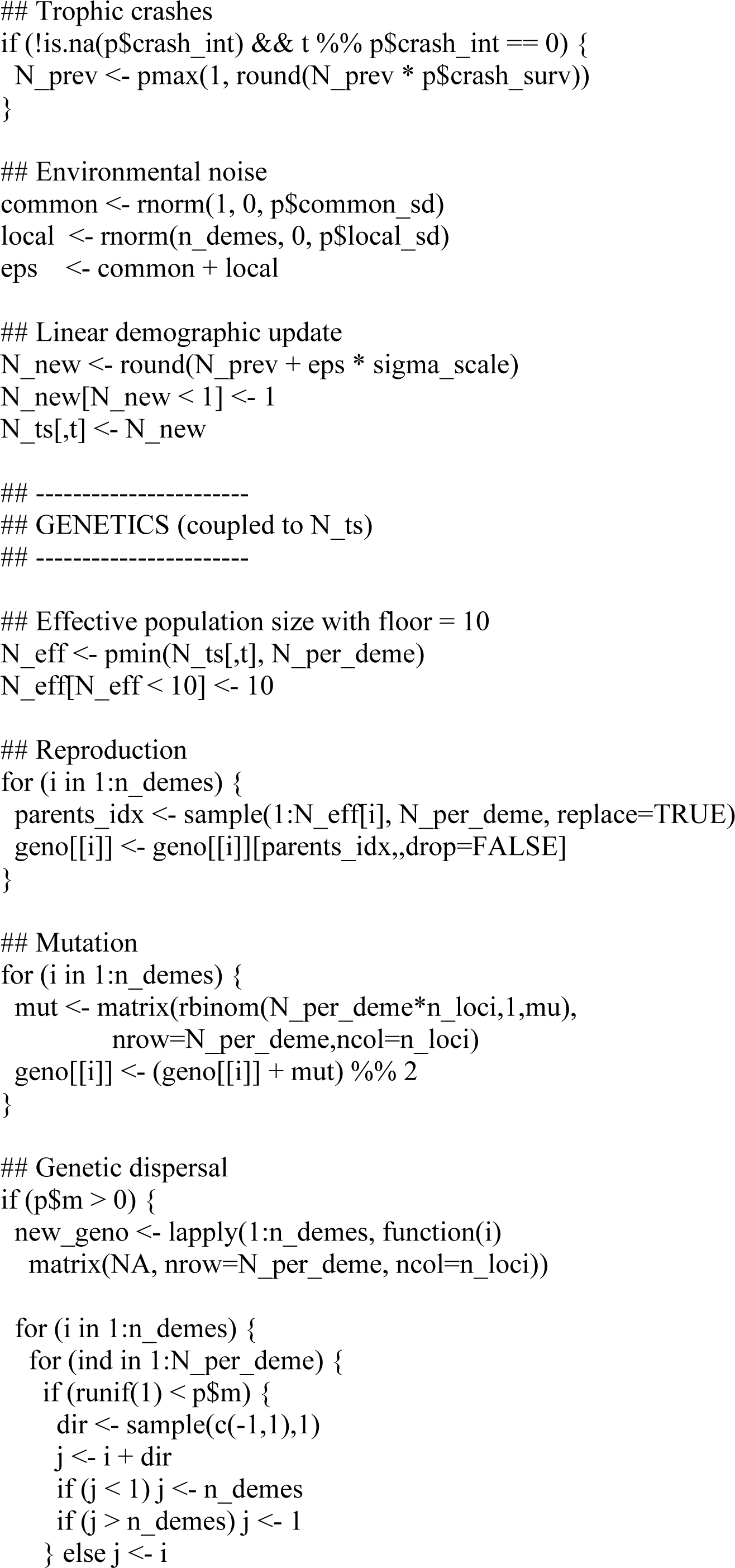
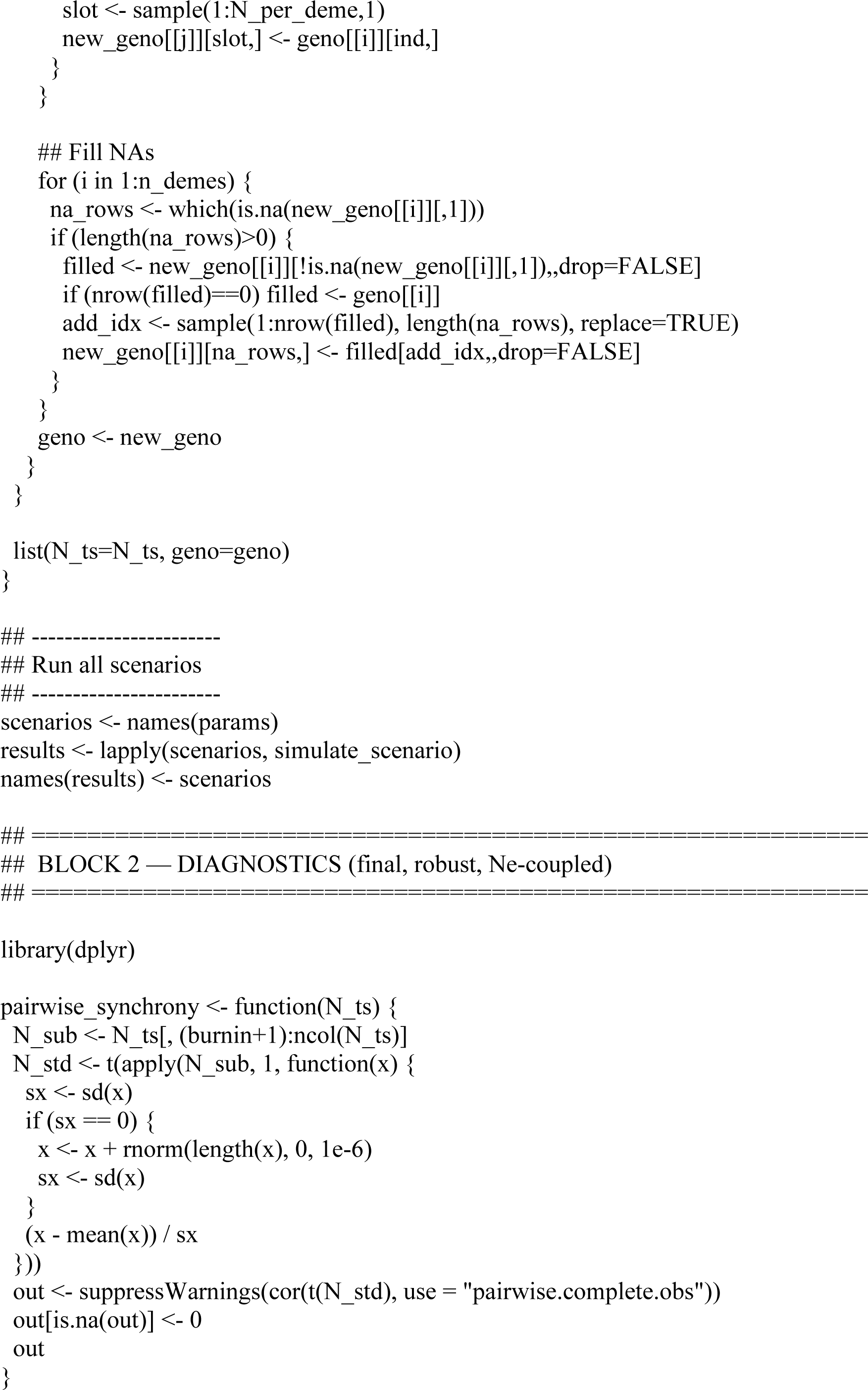
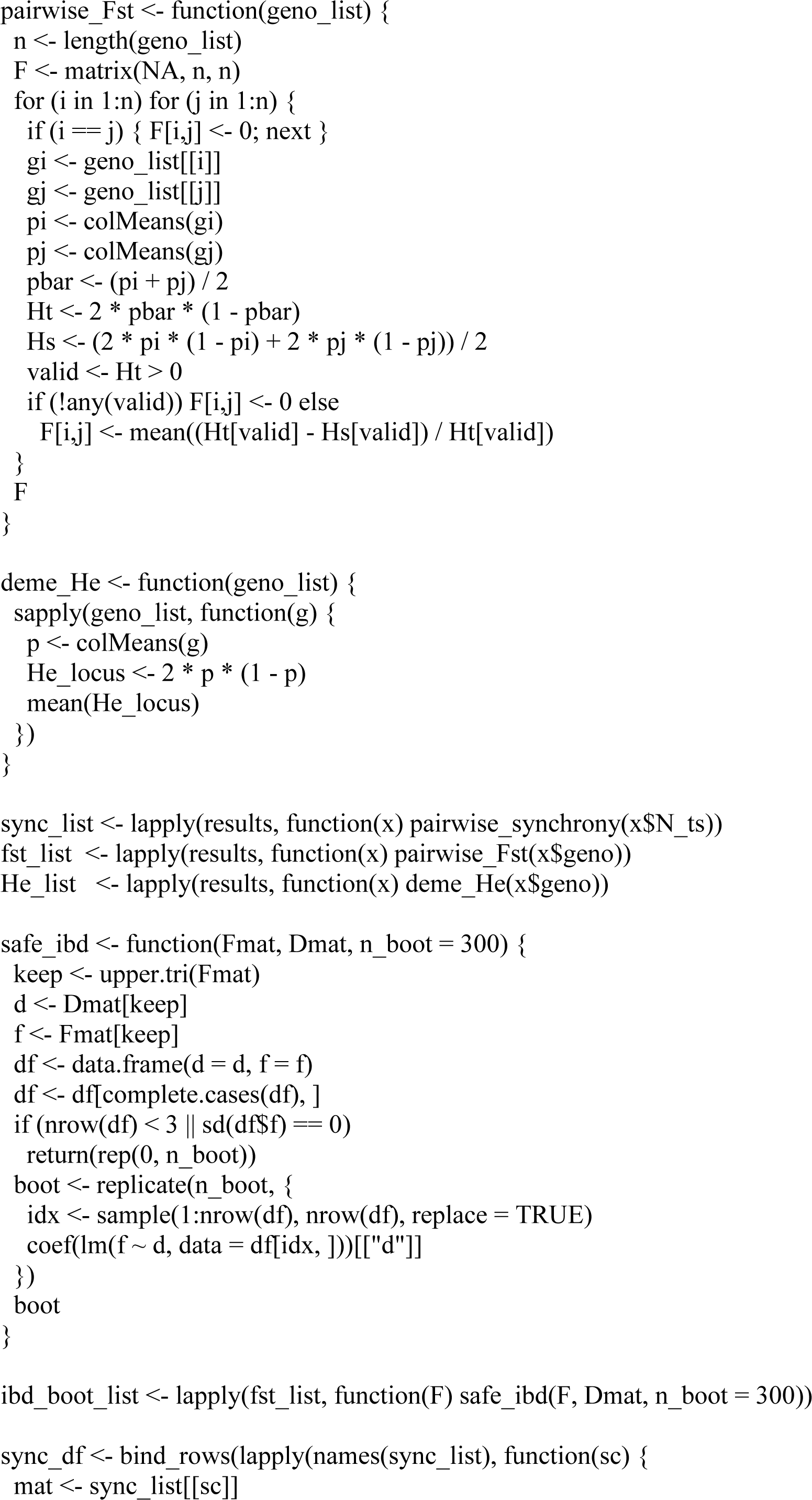
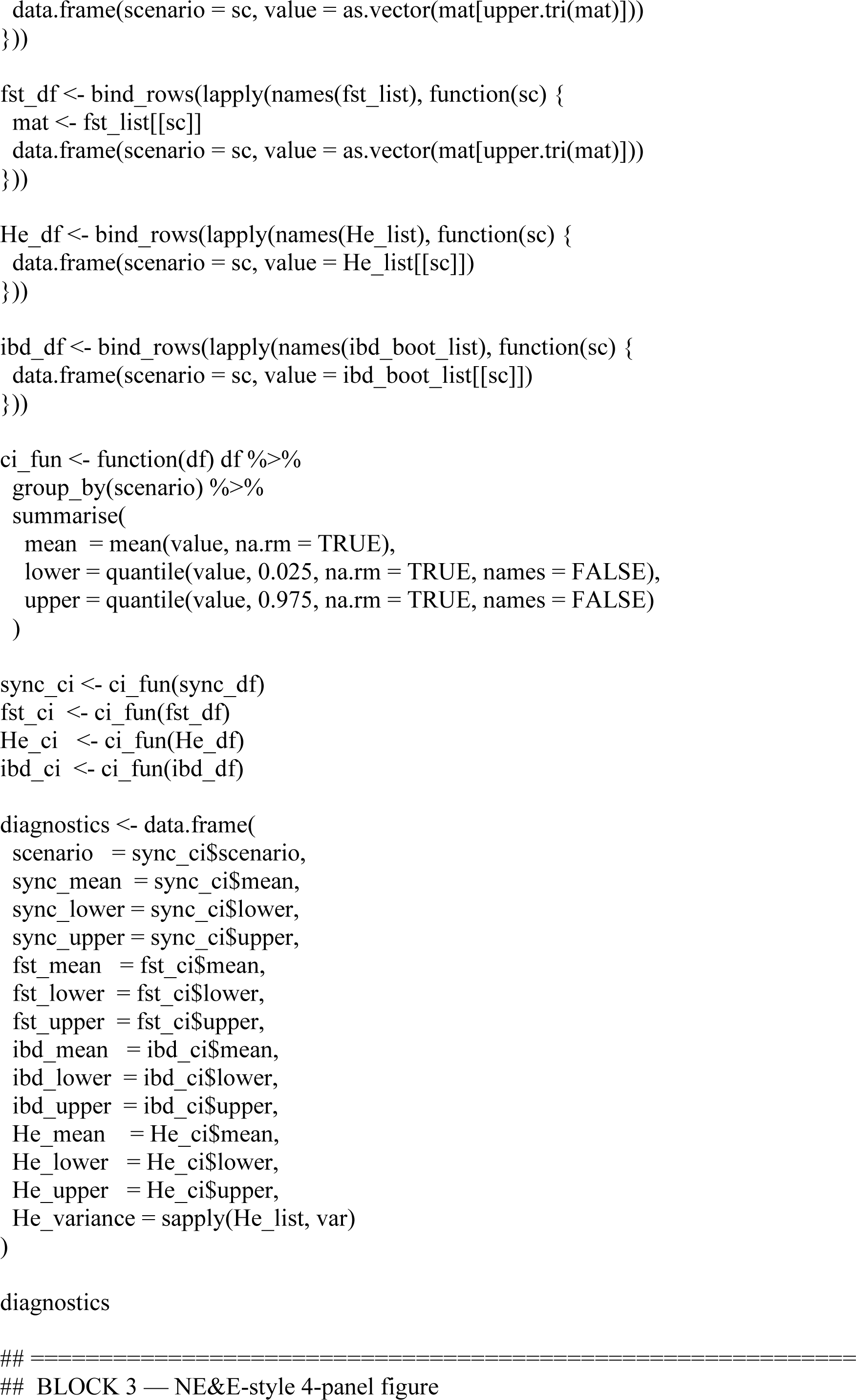
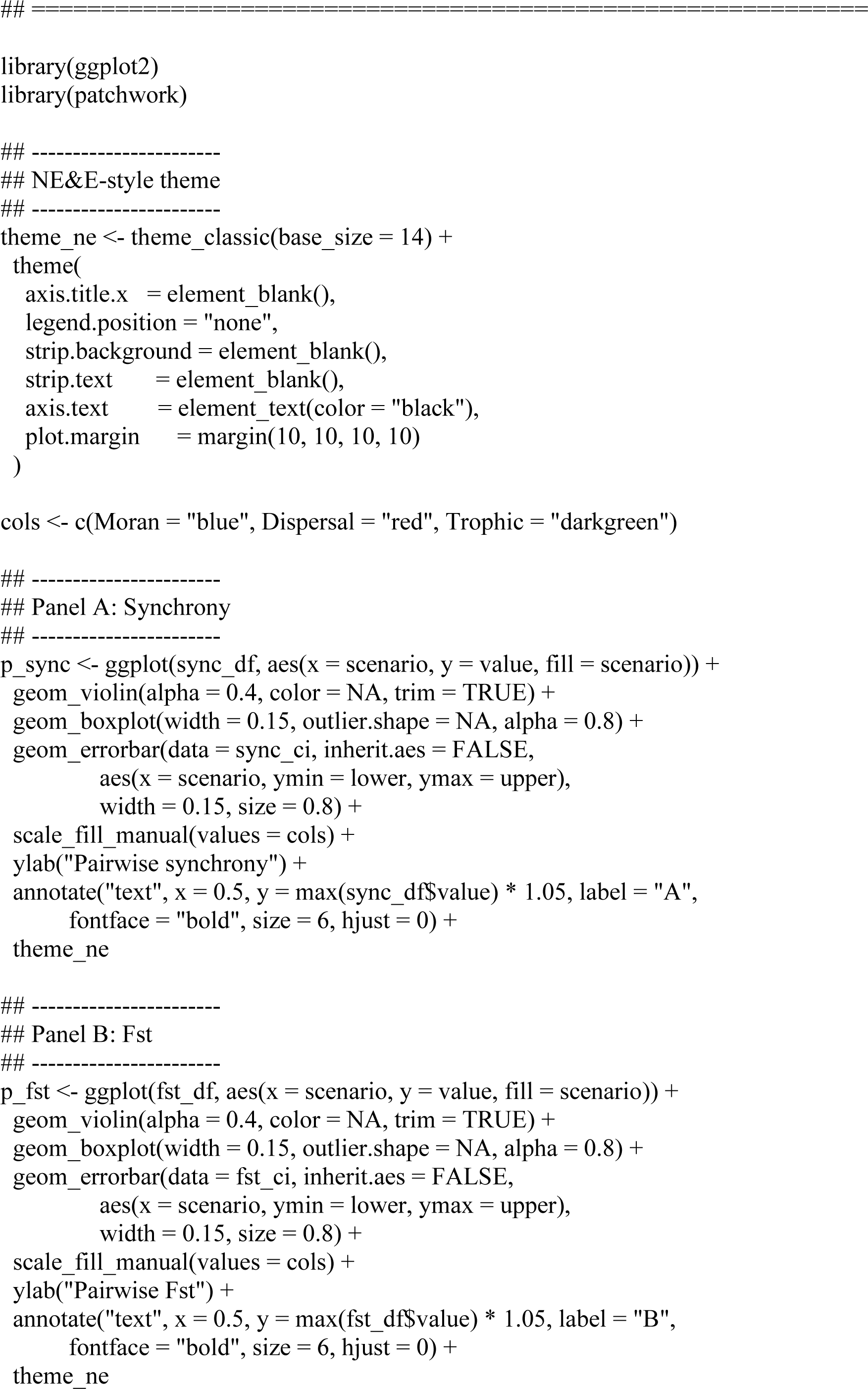
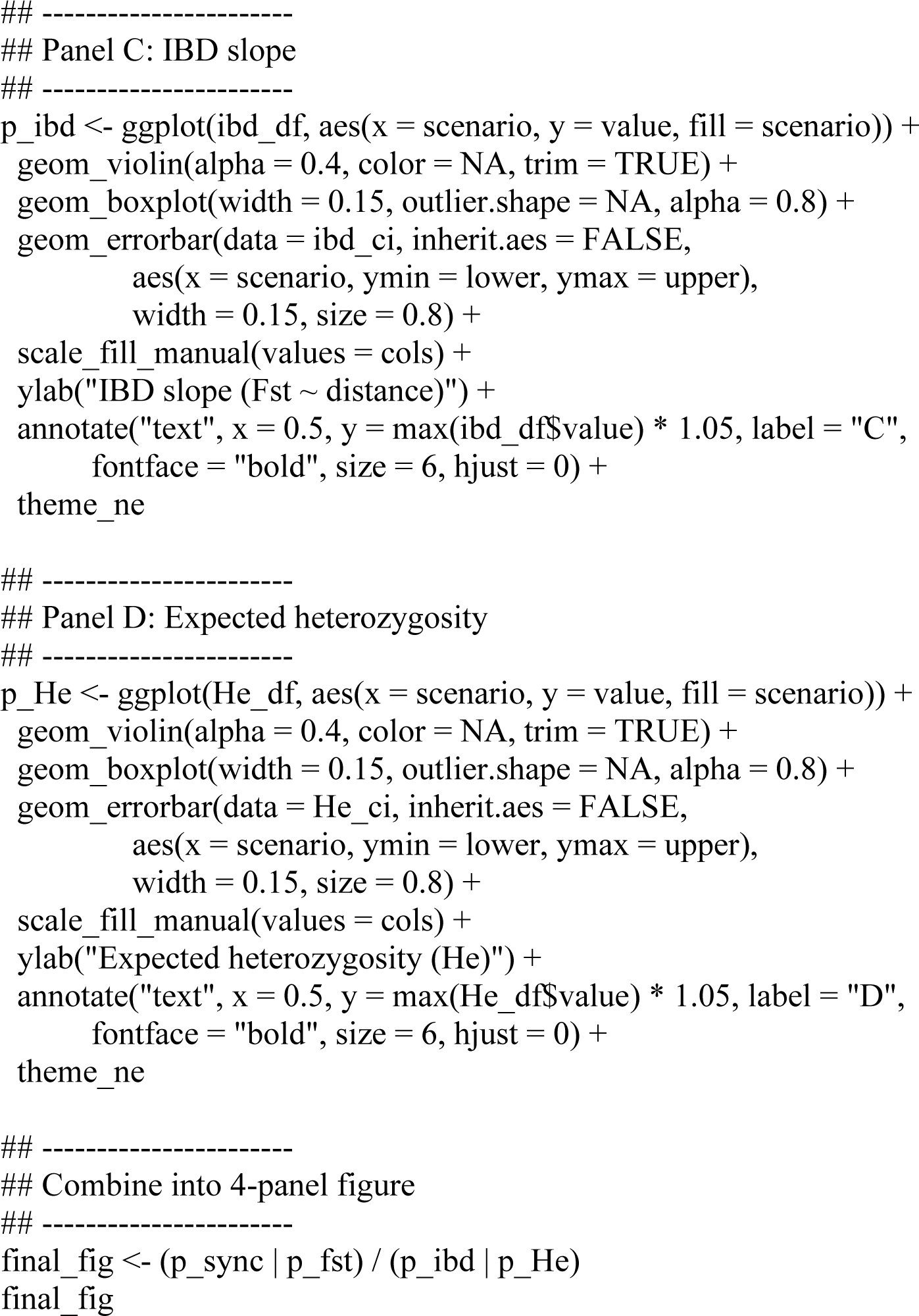

